# Single-Cell Stochastic Modeling of the Action of Antimicrobial Peptides on Bacteria

**DOI:** 10.1101/2021.05.12.443915

**Authors:** Hamid Teimouri, Thao N. Nguyen, Anatoly B. Kolomeisky

## Abstract

Antimicrobial peptides (AMPs) produced by multi-cellular organisms as their immune system’s defense against microbes are actively considered as natural alternatives to conventional antibiotics. Although a substantial progress has been achieved in studying the AMPs, the microscopic mechanisms of their functioning remain not well understood. Here, we develop a new theoretical framework to investigate how the AMPs are able to efficiently neutralize the bacteria. In our minimal theoretical model, the most relevant processes, AMPs entering into and the following inhibition of the single bacterial cell, are described stochastically. Using complementary master equations approaches, all relevant features of bacteria clearance dynamics by AMPs, such as the probability of inhibition and the mean times before the clearance, are explicitly evaluated. It is found that both processes, entering and inhibition, are equally important for the efficient functioning of AMPs. Our theoretical method naturally explains a wide spectrum of efficiencies of existing AMPs and their heterogeneity at the single-cell level. Theoretical calculations are also consistent with existing single-cell measurements. Thus, the presented theoretical approach clarifies some microscopic aspects of the action of AMPs on bacteria.

## Introduction

Antimicrobial peptides (AMPs), which are also called host defence peptides, are essential elements of the innate immune systems in multi-cellular organisms [32]. They generally provide the first line of defence against harmful bacteria without being toxic to the host organisms [4, 10, 15, 32]. Moreover, AMPs support their hosts by exhibiting a variety of critically important biological properties such as anti-viral, anti-fungal, anti-cancer, and anti-inflammatory activities [6, 10, 16, 27]. Due to their antimicrobial actions, AMPs are frequently compared to conventional antibiotics that play a central in modern medicine. However, there are two main differences between these classes of active molecules. First, unlike the typical antibiotics that are composed of single organic compounds, AMPs are made of (mostly positively charged) short sequences of amino acids [15]. Second, despite the fact that bacteria in multi-cellular organisms have encountered antimicrobial peptides for millions of years, acquisition of resistance by a bacterial strain against AMPs is much weaker [7, 10, 15, 31, 32]. These unique characteristics of AMPs stimulated significant efforts in exploring them as alternative therapeutic approaches against pathogenic bacteria and other types of infection [9]. In addition, new applications of AMPs in food production, agriculture and medicine have been also intensively debated [8, 10].

It is widely believed that the antimicrobial peptides kill bacteria via one of two main routes [2, 17]. AMPs might associate to the bacterial cell membrane that leads to the disruption of major processes and causes the formation of pores, allowing for the leakage of essential ions and nutrients and eventually killing the bacterium [4, 17]. Alternatively, absorption of AMPs to the membrane can open new pathways for the peptides to move further inside and to act on various intracellular targets [10, 15]. Recent experiments found that in some systems a large number of peptides (∼10^6^ − 10^7^) is needed to completely saturate the bacterial membrane and to trigger disruptive effects on bacteria [21, 32]. This corresponds to AMP operating at the millimolar/micromolar concentrations. However, there are also AMPs that can kill the bacteria even at much smaller nanomolar concentrations [32]. Such a wide spectrum of activities has been attributed to a variety of possible mechanisms of inhibition [2, 4, 10, 15, 32]. However, the microscopic details of AMPs action on bacteria remain mostly unexplained.

A new direction in exploring the mechanisms of antimicrobial peptides opened a recent quantitative study that investigated the dynamics of AMPs actions by integrating single-cell and population level experiments [25]. A fast absorption and unexpected retention of LL37 antimicrobial peptides in *E*.*coli* bacteria has been observed, which led to a complex heterogeneous behavior in the system. The cell growth was inhibited in one group of cells, while the growing sub-population survived due to the sequestration of AMPs by the non-growing fraction of bacterial cells. One of the striking observations of this study was a heterogeneous growth inhibition even at the single-cell level [25]. But the most important result of this study was a quantitative description of the bacterial inhibition dynamics by AMPs.

The importance of AMPs in protecting the living organisms from various infections and microbes stimulated multiple theoretical investigations [11, 14]. These studies, however, concentrated mostly on the structural aspects of entering the bacterial cell membranes and on the molecular models of pore formation. Since most AMPs are relatively short peptide chains, atomistic and coarse-grained MD simulations have been actively utilized for clarifying the mechanisms of bacterial cell membranes disruption and for developing new potential drugs based on AMPs [11, 14, 29]. However, the dynamic aspects of bacterial inhibition have not been addressed. In addition, current theoretical studies have not considered the effect of stochasticity of underlying biochemical and biophysical processes. At the same time, it has been shown recently that the stochasticity plays an important role in bacterial clearance dynamics by conventional antibiotics [5, 26]. In particular, these investigations led to a new definition of minimal inhibitory concentration (MIC) corrected by stochastic effects. It was suggested that even low concentrations (sub-MIC) could potentially lead to successful clearance of the bacteria [26].

In this paper, we present a new theoretical framework to analyze the bacterial clearance dynamics by AMPs. To take into account the random nature of underlying processes, a discrete-state stochastic approach of bacterial inhibition is developed. In our minimal theoretical model, normal bacterial cell growth, AMPs entering into the cell and stopping the growth are viewed as independent random processes. Using various master equations approaches [13, 18, 26, 28], the inhibition dynamics is explicitly evaluated. Our results suggest that both entering and inhibition are equally important for the efficient functioning of AMPs. Theoretical analysis, which is consistent with available experimental observations for antimicrobial peptide LL37 in *E*.*coli* bacteria, also provides plausible explanations for a large range of AMPs activities and for inhibition heterogeneity at the single-cell level.

## 1 Model

To understand the microscopic mechanisms of inhibition, let us consider a single bacterial cell surrounded initially (at *t* = 0) by *N* antimicrobial peptides as illustrated in Fig. 1. At typical cellular conditions, this number is quite large, *N* ∼10^6^ −10^7^ [21]. The cell can grow and divide into two new cells with a rate *λ*, while the AMPs can enter and attach to the bacterium with a rate constant *a* (Fig. 1). The overall entrance rate is proportional to the number of AMPs outside of the bacterium. The peptides already in the cell can lead to the growth inhibition with a rate constant *k*: see Fig. 1. Here we also assume that the inhibition is an additive process, i.e., the rate of killing the bacterium is proportional to the number of AMPs already inside the cell. We assume that the cell growth rate *λ* is independent of the number of attached peptides because it is normally controlled by other external factors such as the availability of nutrients, temperature, osmotic pressure and many others [19]. The AMPs entrance rate into bacterium depends on the amino acid sequence, the membrane lipid composition, details of peptide-membrane interactions and on the antimicrobial peptide concentration outside of the bacteria [23]. The inhibition rate reflects the biochemical and biophysical processes of membrane disruption that are still not well understood [4, 15]. Note that this is a minimal description of a very complex bacterial clearance by AMPs that takes into account only the most relevant processes.

**Figure 1.**
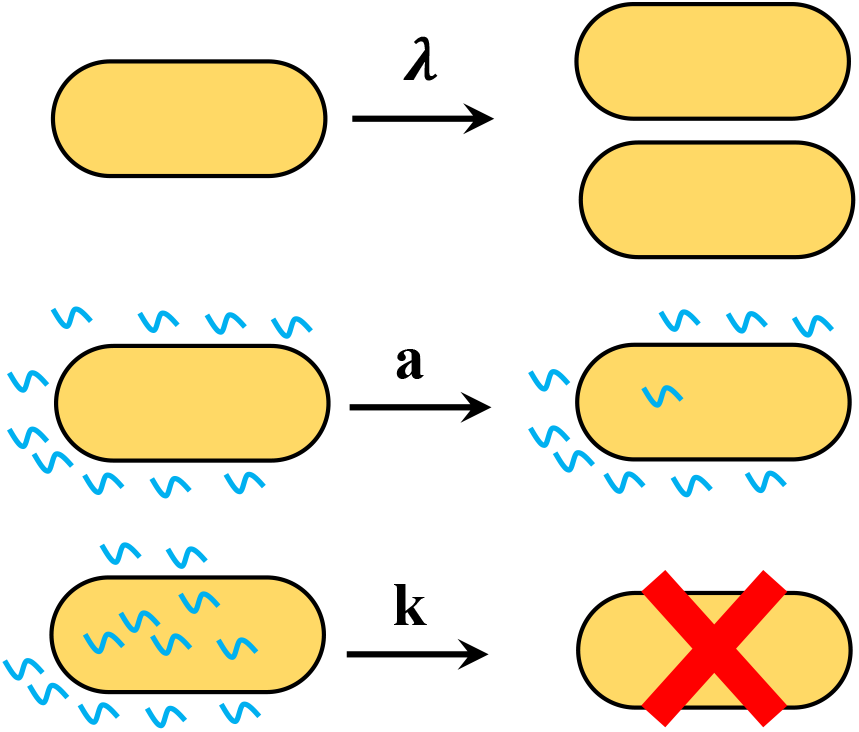
Three fundamental processes that are taking place during the interaction between AMPs and a bacterial cell: growth and division with a rate *λ*, peptides entering the cell with a rate *a* and bacterium inhibition with a rate *k*.

To obtain a comprehensive dynamic description of the system in Fig. 1, we utilize two complementary approaches that are based on master equations exact calculations. We will start with a first-passage probabilities method that has been successfully applied for investigations of various phenomena in Chemistry, Physics and Biology [13, 18]. It concentrates on analyzing the dynamics of first arrival to specific states of the system. The second approach explores conventional forward master equations to evaluate the stationary probabilities of different discrete states in the system [13].

### 1.1 First-Passage Probabilities Analysis

To analyze the system using the first-passage probabilities method, it is convenient to describe the dynamics using a kinetic scheme as presented in Fig. 2. The states are labeled by a variable *n* that describes the number of AMPs already inside the bacterium. From the state *n* (0 *< n < N*) the system can have three outcomes: the cell can grow and divide with the rate *λ*, one more peptide can enter the bacterium with the overall rate (*N* −*n*)*a*, or the cell growth can be inhibited with the overall rate *nk*. In the state *n* = 0 there are no AMPs inside the bacterium and for this reason only the growth (with the rate *λ*) or the peptide entering the cell (with the rate *Na*) are possible. In the state *n* = *N*, all originally available peptides are already in the cell and only the growth (with the rate *λ*) and inhibition (with the rate *Nk*) can happen: see Fig. 2.

**Figure 2.**
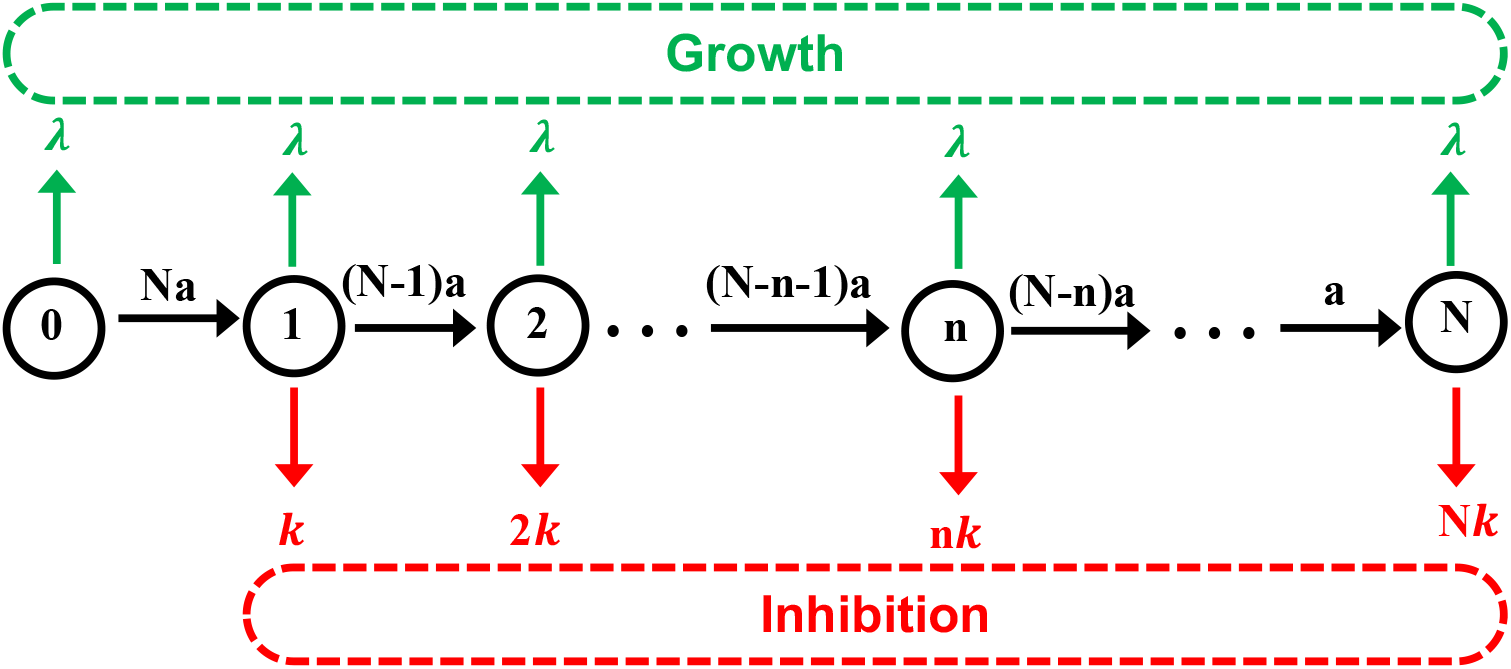
A kinetic scheme for the first-passage probabilities analysis of the bacterial inhibition by AMPs. Each discrete state in the system is labeled as *n* (*n* = 0, 1, …, *N*) where *n* corresponds to the number of already absorbed AMP molecules. Upper (green) arrows describe the uninhibited growth and cell division, lower (red) arrows describe the transitions to the inhibition, and the horizontal (black) arrows correspond to the sequential entrance of antimicrobial peptides into the bacterium. Lower red dashed box corresponds to the fully inhibited state of the bacterium. Upper green dashed box corresponds to the growth state of the bacterium.

The dynamics in the system can be described by defining a function *F*_*n*_(*t*), which is a probability density function for the cell to be inhibited at time *t* given that there are *n* AMPs inside the cell (0 ≤ *n* ≤ *N*) at *t* = 0. The temporal evolution of this function is governed by following backward master equations [13],

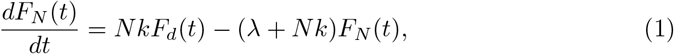

for *n* = *N*, and

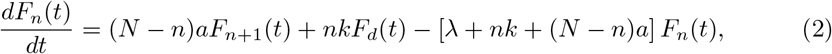

for 0 ≤ *n < N*. Here we also used an additional function *F*_*d*_(*t*) that corresponds to the probability to be inhibited at time *t* for the first time starting already from the inhibited state (see Fig. 2). Clearly, the following boundary condition, *F*_*d*_(*t*) = *δ*(*t*), must be satisfied. The physical meaning of this result is that the bacterial clearance is immediately accomplished if the cell is initially found in the inhibited state.

Introducing the Laplace transforms of the first-passage functions, 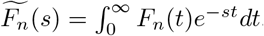, allows us to obtain full exact solutions for backward master equations (see Appendix A.1). Importantly, the first-passage probability density functions contain a comprehensive description of the bacterial clearance process. Specifically, 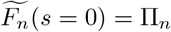 yields the probability of bacterial inhibition given that initially *n* AMP molecules were contained the cell. It can be shown that the inhibition probability Π_*n*_ is given by (see Appendix A.1),

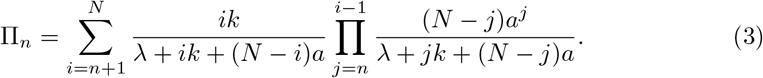

To mimic the experimental observations [25], we are interested in Π_0_ that describes the bacterial clearance probability starting from the state without any peptides inside the bacterium. It can be shown that this probability can be rewritten as,

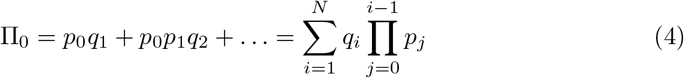

where parameters *p*_*j*_ and *q*_*i*_ are given by

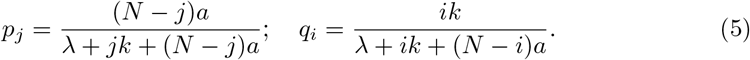

These parameters can be easily understood. The mean residence time at the state *j* is given by 1*/*[*λ* + *jk* + (*N* − *j*)*a*], then *p*_*j*_ is the probability of adding the additional peptide in the state *j* because the corresponding rate is (*N*− *j*)*a*. Similarly, *q*_*i*_ is the probability of inhibition in the state *i* because the corresponding rate is *ik*. These arguments give a clear physical meaning for the expression in Eq. (4). The overall inhibition probability is the sum of inhibition probabilities from each state *i* ≥ 1. The Factor 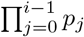 gives the probability for the system to reach the state *i* without bacteria being cleared, and *q*_*i*_ gives the probability of inhibition exactly in the state *i*.

To understand better the bacterial inhibition dynamics, it is interesting to consider two limiting situations. When the entrance rate is much faster than other transitions (*a* ≫ *k, λ*), the system is preferentially found in the state *n* = *N*. This is because from Eq. (5) one can estimate that *p*_*j*_ ≃1, *q*_*j*_ ≃0 for 0 ≤ *j < N*, and *q*_*N*_ = *Nk/*(*λ* + *Nk*). In this limit, the probability of bacterial clearance is given by

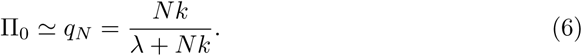

In the opposite limit of very fast inhibition (*k* ≫*a, λ*), the system is is mostly found in the state *n* = 0 because even one peptide inside the bacterium can lead to the inhibition. In this case, we have from Eq. (5) *p*_*j*_ ≃ 0, *q*_*j*_ ≃ 1 for 0 *< j* ≤ *N*, and *p*_0_ = *Na/*(*λ* + *Na*). The probability of bacterial clearance then is given by

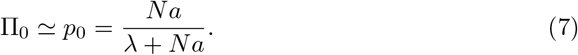

The results of our explicit calculations for the probability of bacterial clearance Π_0_ are presented in Fig. 3. Increasing the entrance and inhibition rates, as expected, leads to higher probabilities of killing the bacterium. But the important result here is that both processes (the entrance and the inhibition) are equally important for the overall inhibition. One can see this from the following arguments. For any specific value of the bacterial clearance probability Π_0_ = Π^*^, from Eqs. (6) and (7) one might conclude that for *a < a*^*^ = *λ*Π^*^*/*[*N* (1 −Π^*^] and/or for *k < k*^*^ = *λ*Π^*^*/*[*N* (1 −Π^*^] it is not possible to achieve the inhibition probability larger than Π^*^. In other words, for any peptide entrance rate below the critical value *a*^*^ it is not possible to reach the inhibition probability above Π^*^ for any possible values of the killing rate *k*. Similarly, for any inhibition rate below the critical value *k*^*^ it is not possible to reach the inhibition probability above Π^*^ for any possible values of the entrance rate *a*. One could also see the important role of the number of AMPs around the bacterium, *N*. Increasing *N* lowers these critical values, i.e., it is easier to kill the bacteria with more available antimicrobial peptides.

**Figure 3.**
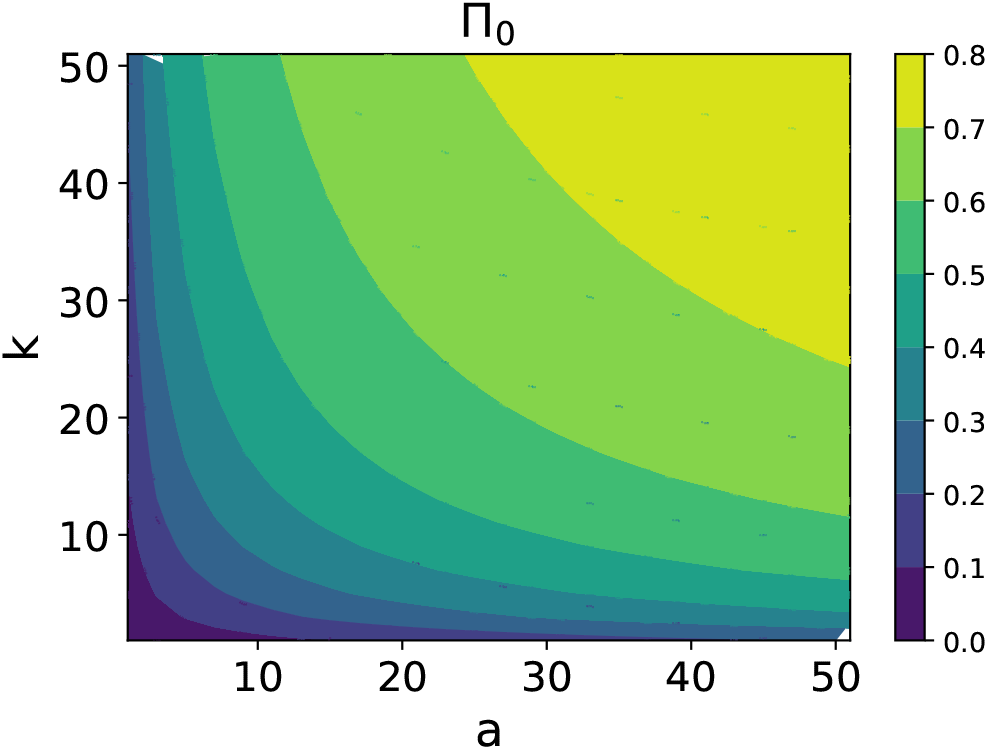
Analytical evaluation of the inhibition probability Π_0_ for a wide range of entrance and inhibition rates *a* and *k* (expressed in units of *λ/N*). For calculations, parameters *N* = 100 and *λ* = 3*/*60 min^−1^ have been used.

Another important characteristics of the bacteria clearance dynamics by AMPs is a mean inhibition time. This time scale corresponds to the average time before the bacteria will stop growing after being exposed to antimicrobial peptides. This property is crucial for developing new AMP-based therapies and for evaluating the bacterial tolerance and resistance [3, 30]. In the language of the first-passage probabilities method, the time *T*_*n*_ corresponds to the mean first-passage time to reach the inhibition state starting initially in the state *n* (see Fig. 2). Using the probability density functions *F*_*n*_(*t*), one can write

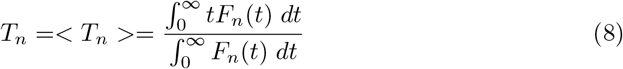

We are mostly interested in the mean inhibition times starting from the state *n* = 0 when there are no AMPs in the bacterium, *T*_0_. As shown in the Appendix A.1, it is given by

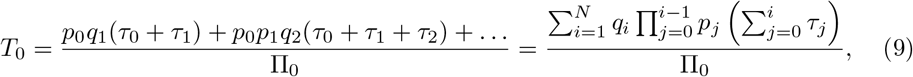

where 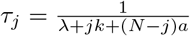 is the residence time for the system to be found in the state *j*.The physical meaning of Eq. (9) is the following. The total mean inhibition time is a sum of *N* contributions corresponding to the bacterial cell killing from the specific state *n* = 1, 2, …, *N*. Each such term is a product of several components. The probability of inhibition at the state *n* = *i* is given by 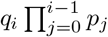, while the second multiplier is the sum of the residence times along this specific inhibition pathway that ends in the state 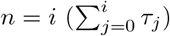. At the end, everything is divided by the total probability of inhibition, Π_0_, to account for those events that do not lead to the bacterial clearance (growth and division with the rate *λ*).

The expression for the mean inhibition time simplifies in the limiting situations. When the peptide entrance rates are very fast (*a* ≫ *k, λ*), the system is mostly found in the state *n* = *N*. The residence times in all states except the state *n* = *N* are very short, *τ*_*i*_ → 0 for *i < N*, while *τ*_*N*_ = 1*/*(*λ* + *Nk*). Then using Eq. (6) we obtain in this limit

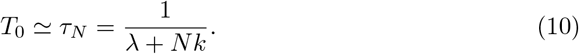

In the opposite limit of very fast inhibition rates (*a* ≫ *k, λ*), the system will not be able to proceed beyond the state *n* = 1, and we have using Eq. (7),

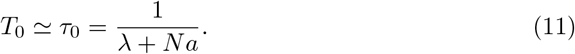

The results of calculations for the mean inhibition times are presented in Fig. 4. Similarly to the inhibition probability, one can see that both entrance and the killing rates are important in order to achieve fast bacterial clearance. For any desired mean inhibition time *T* ^*^, for the entrance and killing rates smaller than 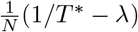 it is not possible to achieve the bacterial inhibition by AMPs at times shorter than *T*^*^. Comparing with Fig. 3, one could also conclude that, as expected, the bacterial clearance probabilities and the mean inhibition times correlate with each other: the larger is Π_0_, the shorter is *T*_0_, i.e., if the probability of inhibition is large it will happen fast.

**Figure 4.**
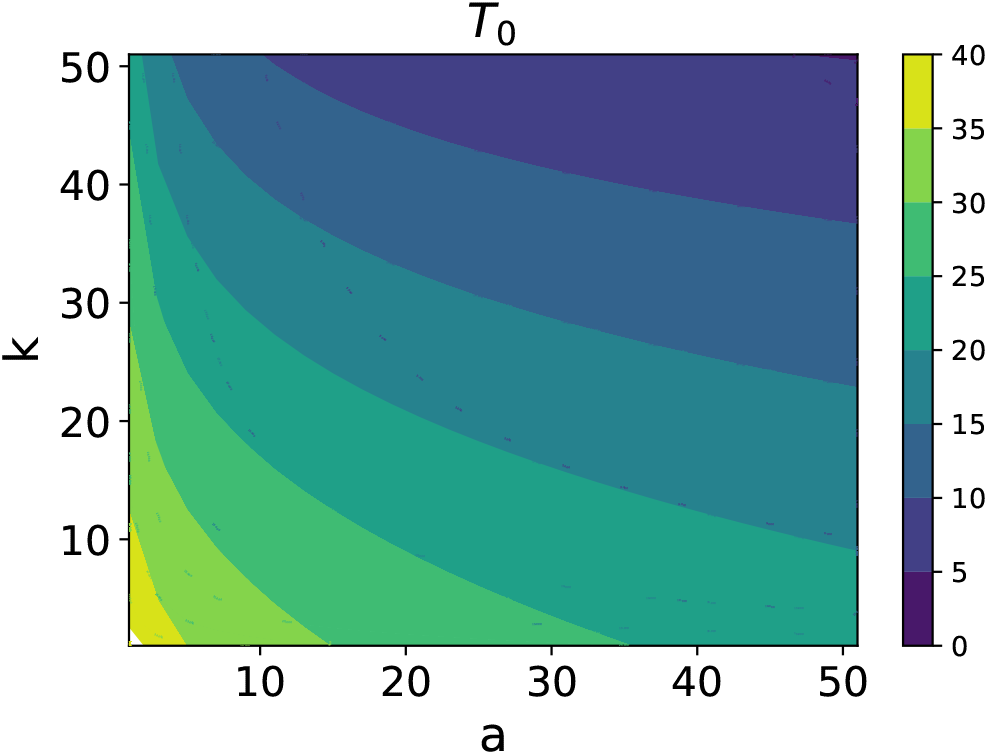
Analytical evaluation of the mean inhibition time *T*_0_ for a wide range of entrance and inhibition rates *a* and *k* (expressed in units of *λ/N*). For calculations, parameters *N* = 100 and *λ* = 3*/*60 min^−1^ were used.

Our theoretical method also allows us also to estimate the degree of fluctuations in the inhibition dynamics by AMPs. It can be done by evaluating the noise, which might be defined as a cell-to-cell variation in the inhibition times. More specifically, we explicitly calculate the normalized variance of inhibition times,

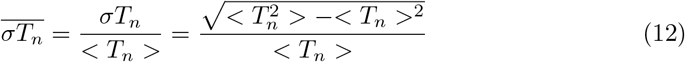

where the expression for second moment 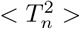 is given in the Appendix A.1. Based on these calculations, one could argue that the level of noise in the system weakly depends on both entrance and killing rates.

### 1.2 Forward Master Equations Approach

There is an alternative method of analyzing the dynamic properties of the AMPs bacterial inhibition by utilizing forward master equations approach. To do so, it is convenient to employ a kinetic scheme shown in Fig. 5. Since we are interested in the inhibition dynamics starting from the situation when there are no AMPs in the bacterium yet, as soon as the system goes to the growth (green transitions) or to the inhibition (red transitions) we reset it back to the state *n* = 0. This explains why all growth and inhibition transitions return to the state *n* = 0: see Fig. 5. Now, one can define a function *P*_*n*_(*t*) as the probability to find the system in the state *n* (which also corresponds to *n* AMPs inside the bacterium) at time *t*. Temporal evolution of these functions is governed by a set of conventional forward master equations,

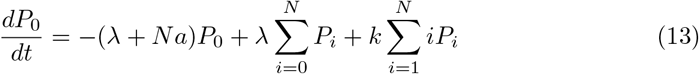

for *n* = 0,

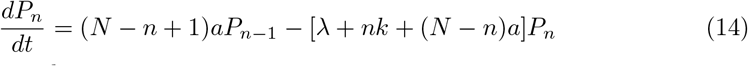

for 1 ≤ *n* ≤ *N* − 1, and

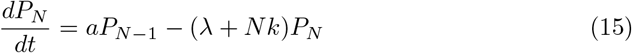

for *n* = *N*. In addition, there is a normalization condition, 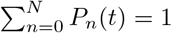.

**Figure 5.**
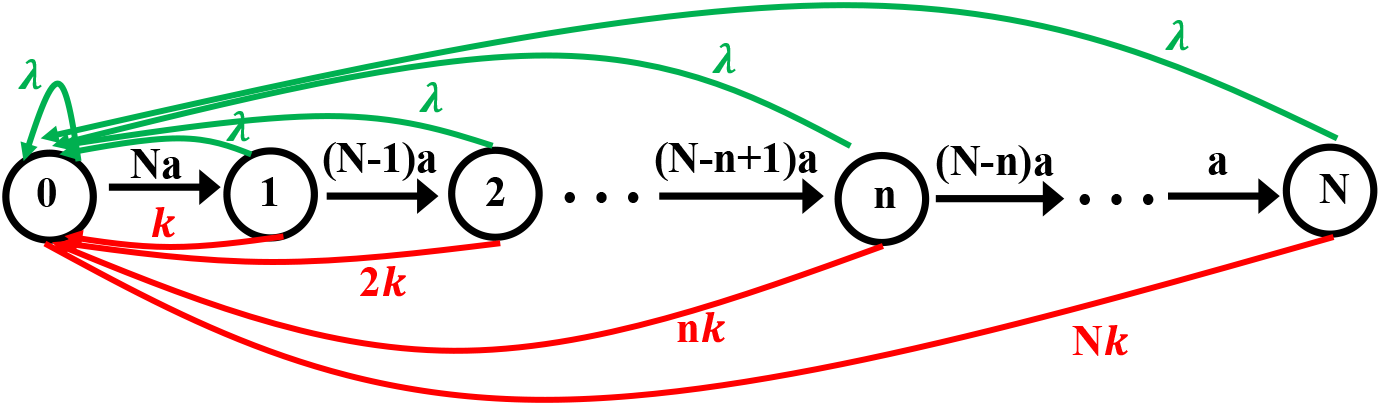
Schematic of the model for the calculation of distribution of number of peptides absorbed by cell.

At long times (*t* → ∞), the system reaches the stationary state where 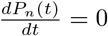, and this allows us to solve explicitly the forward master equations as explained in the Appendix A.2,

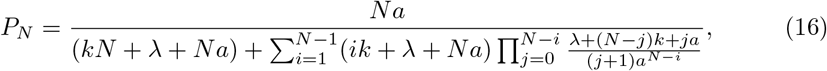

for *n* = *N*, and

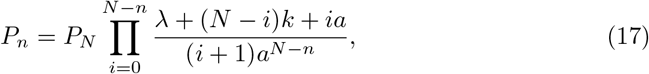

for 1 ≤ *n* ≤ *N* − 1, while

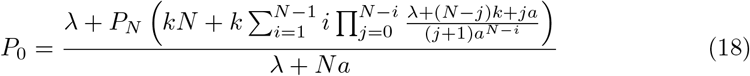

for *n* = 0. It can be easily shown for a special simple case *N* = 1,

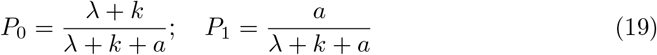

Knowing the stationary probabilities for all states, one can estimate the dynamic properties in the system. The total inhibition flux is given by

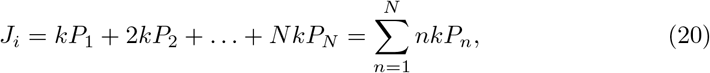

while the total flux in the growth direction is given by

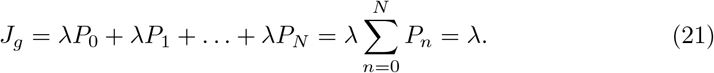

The forward and backward master equations approaches provide a complimentary description of the bacterial clearance dynamics, and the connections between these approaches can be easily established. For example, the inhibition probability calculated explicitly in the first-passage approach can be also estimated using the forward master equations as

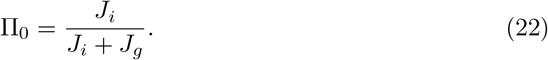

It can be shown explicitly that this expression fully agrees with the one obtained using the first-passage approach: see Eq. (4). The relations between the mean inhibition times and inhibition fluxes are more complex, but they can be provided using the approach discussed in Ref. [1].

The functions *P*_*n*_ not only give the stationary probabilities of different states in the system, but they also provide a stationary distribution of the number of AMPs inside the bacterial cell. One can estimate then the average number of absorbed AMPs needed to clear the bacterial infection,

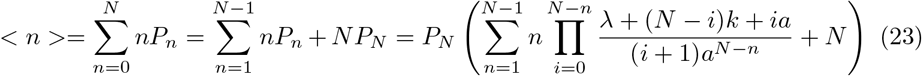

But at the same time, one can rewrite Eq. (20) using the average number of absorbed AMPs, yielding

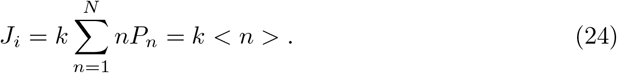

This means that the overall inhibition flux, as expected, is proportional the average number peptides inside the bacterium.

Our theoretical method also allows us to analyze the heterogeneity in inhibition dynamics at the single-cell level by quantifying the variations in the number of absorbed AMPs. For this purpose, we define a dimensionless parameter, known as a Fano factor, which is given by the ratio between the variance and the mean of the number of the absorbed peptides,

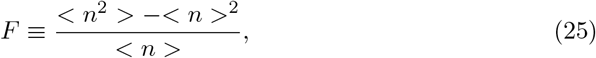

where the second moment in the distribution is equal to

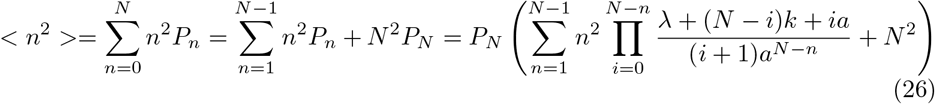

The Fano factor is a convenient measure of heterogeneity because it correlates with the degree of fluctuations in the number of AMPs inside the bacterium.

The results of our explicit calculations for the distributions of absorbed peptides and the Fano factor are presented in Fig. 6. The analysis of the Fano factor (Fig. 6a) shows interesting trends. While increasing the inhibition rate *k* lowers the Fano factor, for larger entrance rates *a* the Fano factor is increasing. This can be explained by analyzing the distributions of absorbed AMPs inside the bacterium. One can see that when the inhibition rate is much faster than the entrance rate the distribution is narrow because the bacterial cell stops growing before the large number of AMPs can enter inside (Fig. 6b). This leads to the smaller fluctuations in the number of peptides inside the bacterial cell. The situation is different for the fast entrance rates (*a* ≫ *k*) when the distribution is wide because many AMPs can enter the cell before it can be inhibited: see Fig. 6c. In this case, the fluctuations are much larger because the inhibition can happen for any number of absorbed peptides. Importantly, these arguments suggest that the Fano factor can be utilized as a quantitative measure of cellular heterogeneity in the inhibition dynamics of AMPs that can also provide the information on the microscopic mechanisms. High heterogeneity would correspond to the systems where the entrance rates are fast, while lower heterogeneity would describe peptides with fast killing rates.

**Figure 6.**
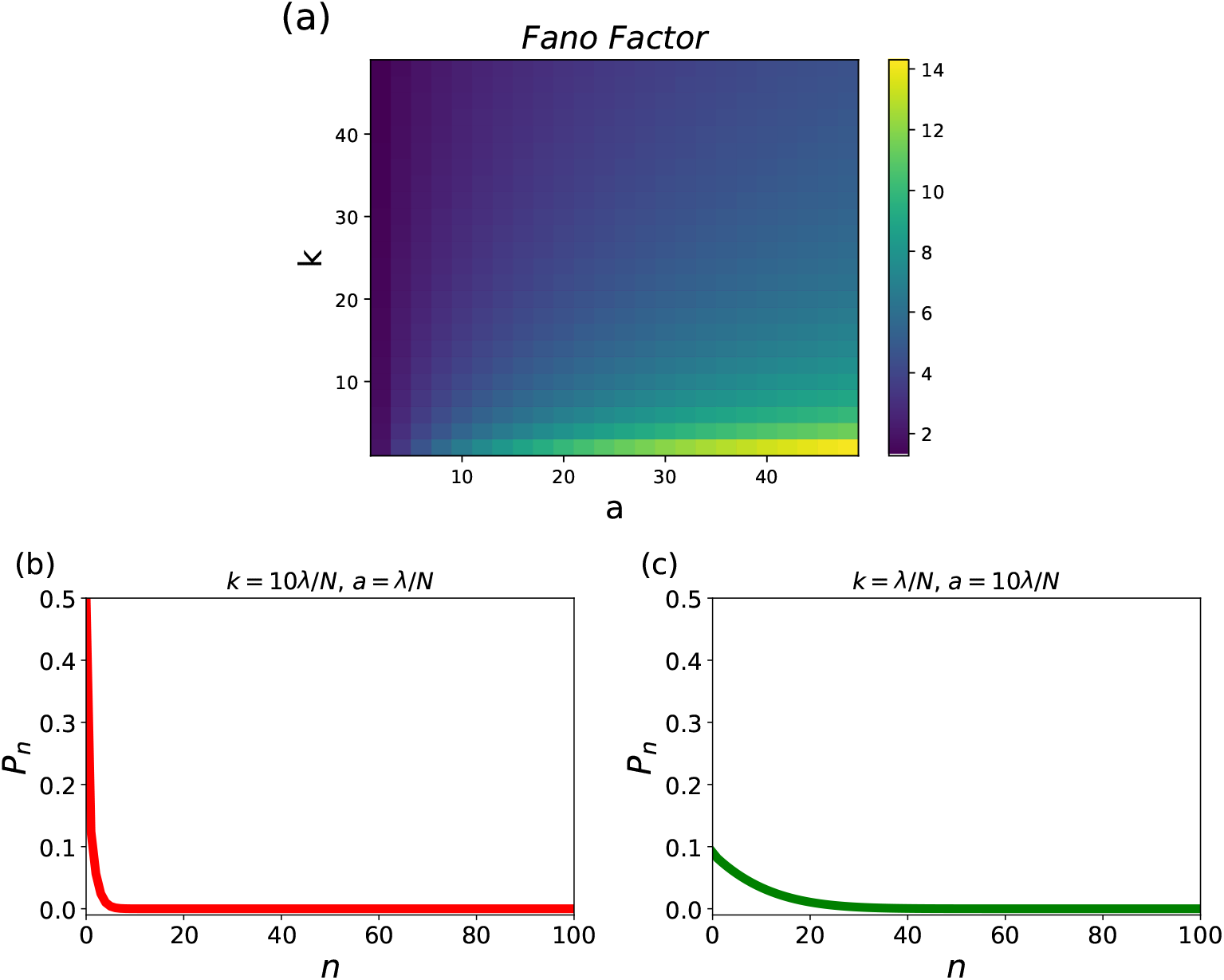
(a) Analytical evaluation of the Fano factor for the number of AMPs inside the bacterial cell for a wide range of entrance and inhibition rates *a* and *k* (expressed in units of *λ/N*). Probability distributions for (b) *k* = 10*λ/N* and *a* = *λ/N* ; and (c) for *k* = *λ/N* and *a* = 10*λ/N*. For calculations, parameters *N* = 100 and *λ* = 3*/*60 *min*^−1^ were used.

To test our theoretical model, it is important to compare its predictions with experimental results. However, there is a limited amount of quantitative information on inhibition dynamics by AMPs at the single-cell level. A recent study provides such data by demonstrating the absorption and retention of LL37 antimicrobial peptides in *E*.*coli* bacteria [25]. The rapid translocation of AMPs followed by the inhibition of bacterial cell growth was monitored using fluorescent time-lapse microscopy that allowed to simultaneously visualize the growth of bacterial cells and the absorbance of AMPs.

While the growth rate for *E*.*coli* bacteria at the conditions utilized in these experiments is more or less is known, *λ* ≃0.03 min^−1^ [25], we need to estimate the entrance rate *a* and the inhibition rate *k*. The following arguments can be presented to approximately evaluate these parameters from experimental data [25]. In experiments, the concentration of AMPs used in single-cell measurements was *c* = 10 *µ*M. Then the volume around the single cell can be roughly estimated as ∼ 100 *µ*m^3^, and there are *N* ∼ 10^6^ peptides per cell in this volume. Experiments suggest that at these conditions in ∼ 10 minutes all AMPs go inside the bacterium. This gives us the estimate of the parameter *a* ∼ 0.1 min^−1^. It seems that the growth does not stop for approximately 10-20 minutes, allowing us also to approximate *k* ≃ 10^−7^ min^−1^. Since the entrance rate *a* is much faster than other transitions in the system, we can utilize Eqs. (6) and (10) to estimate the probability of inhibition and the mean inhibition time,

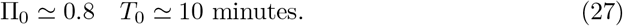

Note that although our estimates of the transition rates in the system are quite crude, our predictions for the mean inhibition times are consistent with experimental observations [25]. This gives a support to our theoretical model.

These calculations also suggest that our theoretical framework might naturally explain a very wide range of concentrations at which different AMPs are functioning. We can argue that those peptides that operate in the millimolar/micromolar range probably have faster entrance rates, *a > k* so that a significant number of peptides is needed to enter the bacterium in order for the inhibition to be achieved. At the same time, AMPs working in the nanomolar regime probably have faster inhibition rates (*k > a*) since even a relatively small number of absorbed peptides can stop the growth of the bacterium cell. It seems that LL37 peptides are operating in the *a > k* regime that corresponds to larger heterogeneity in the system.

We developed a new theoretical framework for analyzing the bacterial clearance dynamics by AMPs at the single-cell level. This is done by taking into account the most relevant stochastic processes, such as the bacterial growth, AMPs entrance and inhibition, as random stochastic processes. Using two complementary master equations approaches, the dynamic properties of bacterial inhibition are explicitly evaluated. It is found that both the entrance and the killing processes are equally important to support the effective action of AMPs. The probability of inhibition correlates with the speed of bacterial clearance. It is also shown that our theoretical predictions are consistent with available experimental data. In addition, it is suggested that the proposed theoretical approach might explain the wide spectrum of activities of various AMPs that operate at different concentration ranges. Furthermore, we presented a specific quantitative measure of heterogeneity and explained that larger heterogeneity is associated with faster entrance rates, while the faster inhibition rates do lead to a smaller heterogeneity.

Since the development of new AMP-based therapeutics is critically important for multiple medical applications, our theoretical results might help in designing new AMPs. We suggest that one should concentrate not only on improving the killing abilities of peptides when they are already bound to bacteria, but also on the way for AMPs to enter faster. In addition, by tuning the entrance and the killing rates one might control the heterogeneity in the action of AMPs that be also important for applications. It will be interesting to test in experiments this combined strategy.

Although the presented theoretical framework is able to provide a comprehensive description for the inhibition dynamics of antimicrobial peptides on bacteria, it should be emphasized that the model is very simplified and several important aspects of the system need to be explored more. Clearly, the processes of entering into the cell and the inhibition involve multiple biochemical and biophysical transformations and cannot be well approximated as single one-step stochastic transitions. In addition, it is important to consider various bacterial strategies of resistance to AMPs [12, 20]. This includes the sequestration by bacterial surface structures, the alteration of membrane charges and/or fluidity, the degradation and removal of AMPs by the efflux pumps. Also bacteria may exhibit collective tolerance effects through variety of mechanisms, including the membrane-displayed proteases that degrade AMPs directly [22]. It will be interesting to extend and generalize our theoretical approach in these important directions [20, 22, 24, 30].

## Acknowledgments

The work was supported by the Welch Foundation (C-1559), by the NSF (CHE-1953453 and MCB-1941106), and by the Center for Theoretical Biological Physics sponsored by the NSF (PHY-2019745).

## Author contributions

H.T. and A.B.K designed the research; H.T. and T.N.N. performed the research; H.T. and A.B.K wrote the manuscript.

## Appendix

### A.1. Exact Solutions for the First-Passage Approach

In this appendix, we present the details of calculations for the bacterial inhibition dynamics by AMPs. As shown in the main text, the temporal evolution of the first-passage probability function is governed by following backward master equations [18],

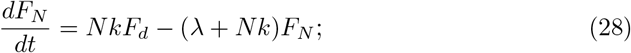

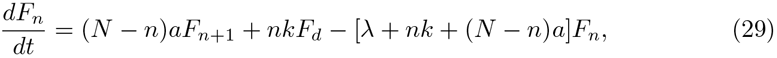

with the initial condition,

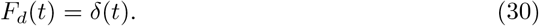

Introducing the Laplace transforms, 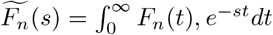 we obtain,

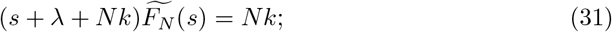

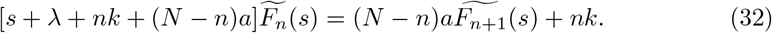

Defining 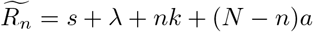, Eqs. (31) and (32) take a simpler form,

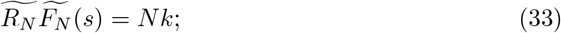

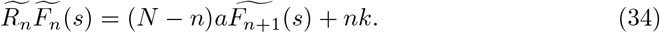

Solving these equations yields,

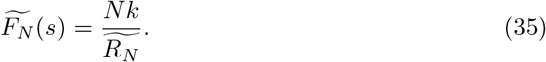

and,

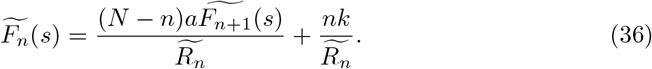

Similarly, for *n* = *N* − 1, we obtain,

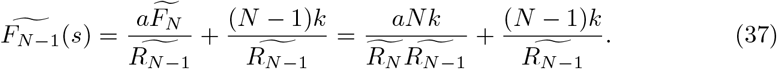

And for *n* = *N* − 2 we have

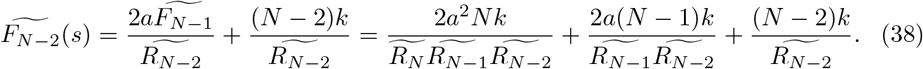

Thus, the general expression for 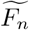 can be presented as

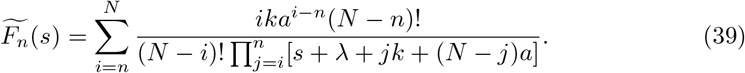

It can be rewritten as

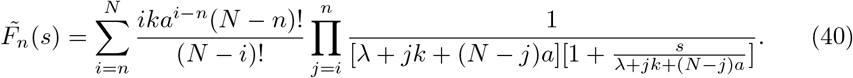

The Taylor expansion of Eq. (40) gives the inhibition probability Π_*n*_:

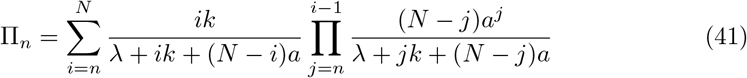

To calculate the mean-first passage times, we need to compute the first derivative of 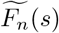. To do so, we rewrite 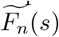 as follows,

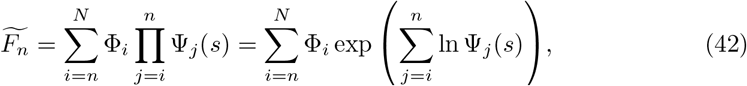

where 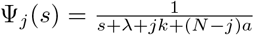 and 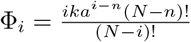 Thus, the derivative of 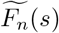 is given by

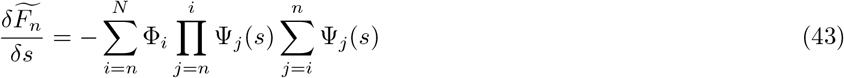

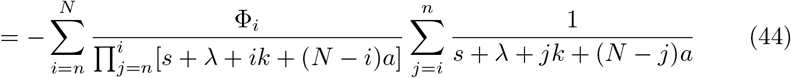

One can now calculate the mean-first passage times from following expression,

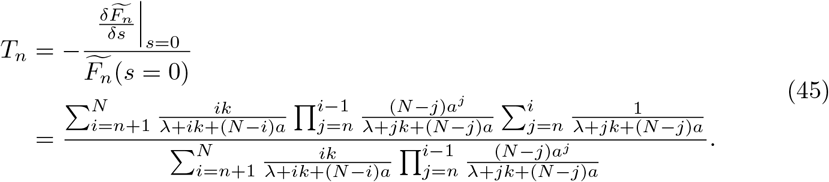

The normalized variance of the first passage times is given by,

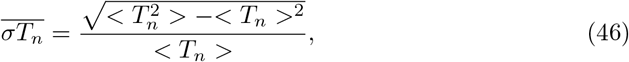

where the second moment is

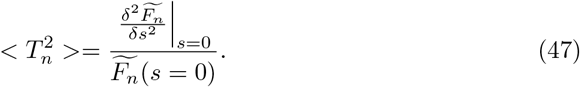

The second derivative of 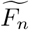 can be easily calculated from Eq. (42), producing

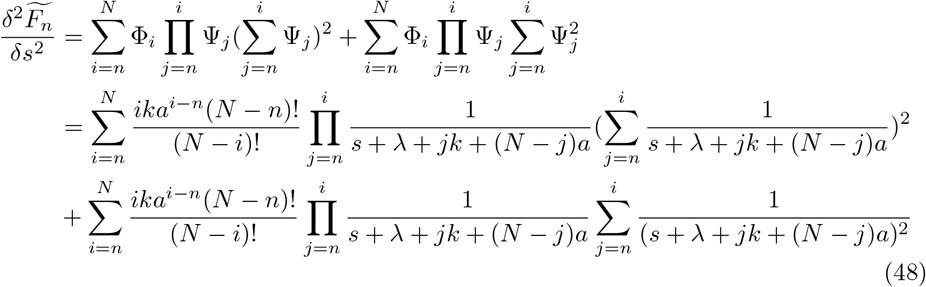

Thus, the second moment is given by

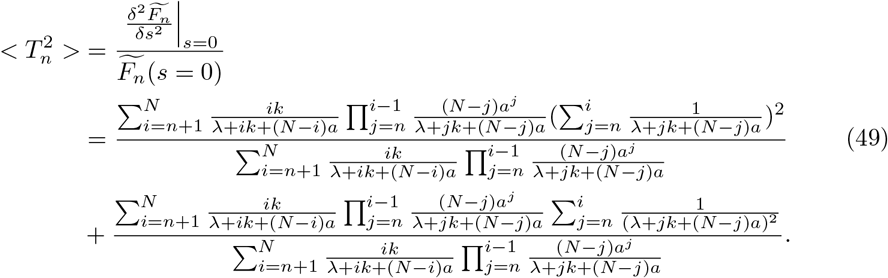

In Fig. 7, we plotted the normalized variance as a function of the transition rates *a* and *k*. One can see a weak dependence on the entrance and inhibition rates.

**Figure 7.**
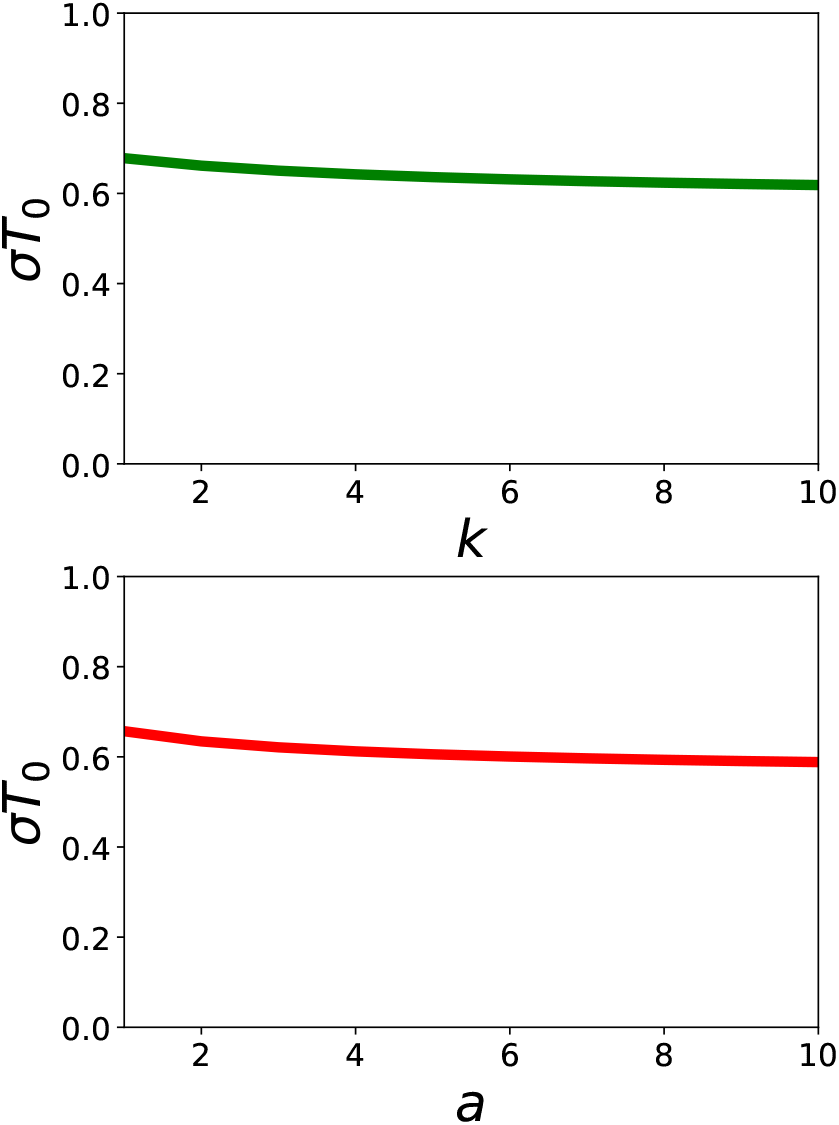
Variance of inhibition times as a function of the translocation rate *a* and the inhibition rate *k*. For calculations, *N* = 100 and *λ* = 0.05 min^−1^ are used.

### A.2. Calculations Using Forward Master Equations

In this section, we present detailed calculations for dynamic properties of the bacterial clearance system. It is now described by the scheme presented in Fig. 5. The dynamical evolution of the system is governed by the following forward master equations,

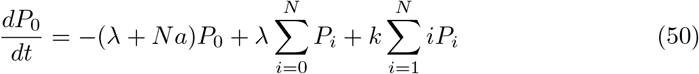

for *n* = 0,

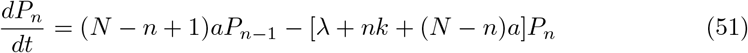

for 1 ≤ *n* ≤ *N* − 1,and

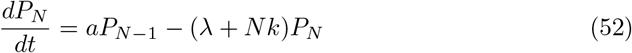

for *n* = *N*. At long times *t* → ∞, the left side of Eqs. (50) - (**??**) becomes equal to zero. After some algebra, we obtain

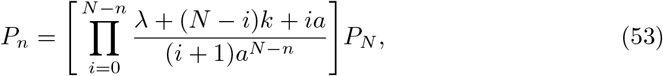

where 1 ≤*n* ≤*N* − 1. One can calculate the constant *P*_*N*_ by substituting Eq. (53) into following expression,

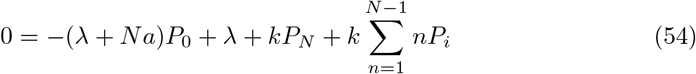

This leads to

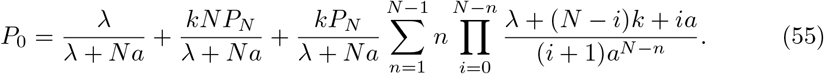

Combining Eq. (55) with the normalization condition 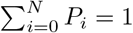, we get

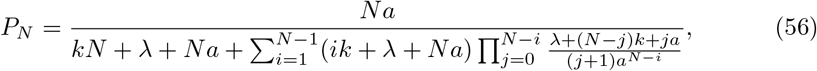

and

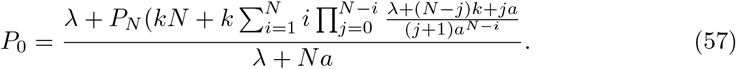

Now the average number of absorbed AMPs can be estimated as

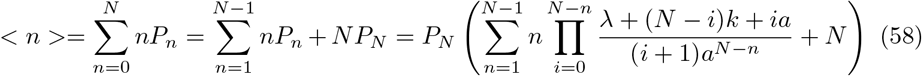

and, the second moment *< n*^2^ *>* is given by

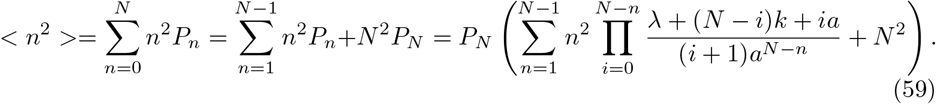

From these equations, we can finally calculate the Fano factor,

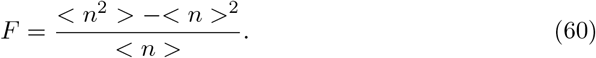

